# A Fast Lasso-Based Method for Inferring Higher-Order Interactions

**DOI:** 10.1101/2021.12.13.471844

**Authors:** Kieran Elmes, Astra Heywood, Zhiyi Huang, Alex Gavryushkin

## Abstract

Large-scale genotype-phenotype screens provide a wealth of data for identifying molecular alterations associated with a phenotype. Epistatic effects play an important role in such association studies. For example, siRNA perturbation screens can be used to identify combinatorial gene-silencing effects. In bacteria, epistasis has practical consequences in determining antimicrobial resistance as the genetic background of a strain plays an important role in determining resistance. Recently developed tools scale to human exome-wide screens for pairwise interactions, but none to date have included the possibility of three-way interactions. Expanding upon recent state-of-the art methods, we make a number of improvements to the performance on large-scale data, making consideration of three-way interactions possible. We demonstrate our proposed method, `Pint`, on both simulated and real data sets, including antibiotic resistance testing and siRNA perturbation screens. `Pint` outperforms known methods in simulated data, and identifies a number of biologically plausible gene effects in both the antibiotic and siRNA models. For example, we have identified a combination of known tumor suppressor genes that is predicted (using `Pint`) to cause a significant increase in cell proliferation.

**Author Summary:** In recent years, large-scale genetic datasets have become available for analysis. These large datasets often stretch the limits of classic computational methods, requiring too much memory or simply taking a prohibitively long time to run. Due to the enormous number of potential interactions, each gene or variation in the data is often modeled on its own, without considering interactions between them. Recently, methods have been developed to solve regression problems that include these interacting effects. Even the fastest of these cannot include threeway interactions, however. We improve upon one such method, developing an approach that is significantly faster than the current state of the art. Moreover, our method scales to three-way interactions among thousands of genes, while avoiding a number of the limitations of previous approaches. We analyse large-scale simulated data, antibiotic resistance, and gene-silencing datasets to demonstrate the accuracy and performance of our approach.

## 1. Introduction

Epistatic gene interactions have practical implications for personalised medicine, and synthetic lethal interactions in particular can be used in cancer treatment [4]. Discovering these interactions is currently challenging at a practical scale [35, 25, 14, 16], however. In particular, there are no methods able to infer three-way effects. For a given number of genes there are exponentially many potential interactions, complicating computational methods. If we restrict our attention to pairwise effects, it is possible to experimentally knock out particular combinations of genes to determine their combined effect [13]. This approach does not scale to the approximately 200 million pairwise combinations possible among human protein coding genes, however, let alone 1.3 trillion three-way combinations. We instead consider inferring interactions from large-scale genotype-phenotype data. These include mass knockdown screens, in which a large number of genes are simultaneously suppressed, and the resulting phenotypic effects are measured.

We have shown in [16] that a lasso-based approach to inferring interactions from an siRNA perturbation matrix is a feasible method for large-scale pairwise interaction detection. In this additive model, we assume fitness is a linear combination of the effects of each gene, and the effect of every combination of these genes. For the sake of scalability, in [16] we considered only individual and pairwise effects, and assumed gene suppression was strictly binary. The fitness difference (compared to no knockdowns) in each experiment is then the sum of individual and pairwise effects 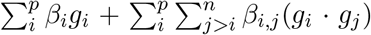, where *g_i_* = 1 if gene *i* is knocked down, 0 otherwise. With sufficiently many such mass-knockdowns, we can infer pairwise interactions by finding the pairs of genes whose effect is not the sum of the effects of each gene individually.

Neither of the previously tested inference methods for this model, `glinternet` and `xyz`, are effective at the genome-scale however. `glinternet` suffers from prohibitively long running times,^1^ and `xyz` does not accurately predict effects in our larger simulations. Our aim is to fit a model including all *p* ≊ 20, 000 human protein-coding genes, with as many as *n* = 200, 000 siRNAs. Furthermore, we aim to go beyond pairwise interactions to consider three-way effects.

A recently developed method, WHInter [39], is effective at solving lasso regression on much larger scale data than `glinternet`. This performance comes as a result of pruning the interactions considered based on the current regularisation parameter, removing interactions that could not possibly have a non-zero effect. Because doing so does not affect the solution of the regression problem, we expect comparable accuracy from WHInter and `glinternet`. Nonetheless there are a number of areas in which we can improve upon WHInter. In particular WHInter does not make use of multi-core CPUs, and considers only pairwise interactions.

We have developed an R-package, `Pint`, that is able to perform square-root lasso regression on all pairwise interactions on a one thousand gene screen, using ten thousand siRNAs, in 15 seconds, and all three-way interactions on the same set in under ten minutes. We are also able to find the largest 2, 000 effects, excluding three-way combinations, on a genome-scale data set with 19,000 genes and 6, 700 siRNAs in half an hour using two eight-core CPUs. This is made possible by taking into account that our input matrix *X* is both sparse and strictly binary, parallelising the pruning method from [39], and compressing the active set. To allow threeway interactions, we extend to a two-step pruning method able to rule out both pairwise and three-way interactions. Our package, `Pint`, is available at github.com/bioDS/pint.

Our method is based on an existing fast algorithm [61], which we adapt for use on binary matrices. We further add parallelised version of the pruning step from [39], expanded to include three-way interactions. We provide a detailed description of our implementation, followed by the scalability analysis, below. We also perform a simulation study to compare our method’s scalability with known methods, and analyze two large-scale experimental data sets.

In the first, an siRNA perturbation screen from [46], we search for both individual genes and combinations (pairwise or three-way) that have an effect when simultaneously silenced, stopping after the first 131 effects have been identified. The results include 22 individual, 41 pairwise and 68 three-way effects. We investigate the biological plausibility of the top five effects, and find that three out of five are suppressing genes that could be related to cell survival. One combination in particular involves simultaneously disabling two tumor suppressing genes, and is predicted to cause a significant increase in cell proliferation.

The second data set is composed of genetic variants identified in the intrinsically antibiotic resistant bacteria *Pseudomonas aeruginosa*. *P. aeruginosa* is an opportunistic pathogen found in a variety of environments and is a leading cause of morbidity and mortality in immunocompromised individuals or those with cystic fibrosis [23, 37]. *P. aeruginosa* is known to acquire adaptive antibiotic resistance in response to long term usage of antibiotics associated with chronic infections [10, 43, 44]. The genomes included in that data set are from strains that have been isolated from chronic and acute infections as well as environmental samples. The minimum inhibitory concentration for the antibiotic Ciprofloxacin has been used as the phenotypic marker for this dataset. Ciprofloxacin belongs to the fluoroquinolone class of bactericidal antibiotics that targets DNA replication and is one of the most widely used antibiotics against *P. aeruginosa* [50].

This set contains over 170,000 SNVs, too many for our method to include all possible three-way interactions. Three-way interactions can be included if we remove columns with less than 30 entries, reducing the matrix to ≈ 76,000 columns. This ignores over half of the SNVs present however, so we instead limit the search to individual and pairwise combinations of variants, and determine the first 50 effects that are discovered. Among these, 13 of the 16 non-synonymous variants were identified as having possible contributions to Ciprofloxacin resistance.

## 2. Materials and Methods

Throughout the paper we refer to fitness landscapes, and focus on fitness values as our phenotype of interest, but would like to note that the theory can be applied for any (binary) genotype-phenotype mapping and any phenotype. Let a fitness landscape be a mapping 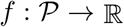 from the genotype space to fitness values. Furthermore, suppose genotypes are strictly binary, 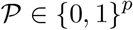, where 1 indicates the presence of a particular mutation (or variant), 0 indicates its absence, and *p* is the number of genes. The complete fitness landscape then describes the effects of all combinations of mutations [5]. For example, the two gene space 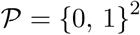 contains the wild-type 00, two single mutants 01 and 10, and the double mutant 11. The fitness landscape *f*: {0,1}^2^ → ℝ in this case can be written as

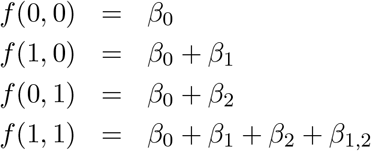

for parameters *β_i_* ∈ ℝ. *β*_0_ is called the bias, *β*_1_ and *β*_2_ main effects, and *β*_1,2_ the pairwise interaction. This pairwise interaction is exactly the classic definition of epistasis in quantitative genetics [17]. In generalising to higher-order interactions, we follow the definitions of Otwinowski and Nemenman [42]. For *p* ≥ 3 genes, the complete fitness landscape *f* is:

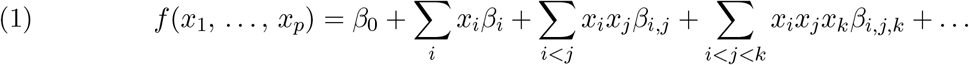

While including all possible interactions quickly becomes computationally and statistically intractable for large *p*, we can model the interactions up to a point. Ignoring interactions of order four and higher we obtain an approximation of the fitness landscape:

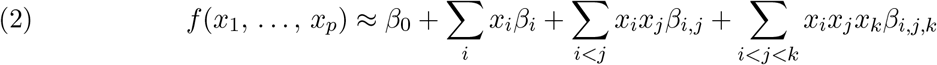

The remainder of this section describes the regression model we use to estimate these effects, the algorithms we use to efficiently solve it, and finally the data sets on which we apply it.

### 2.1. Regression Model

Given as input a matrix **X** ∈ {0,1}^*n*×*p*^ and a vector **Y** ∈ **R**^*n*^, where columns of **X** are genes or variants of interest, rows are samples from the genotype space, and entries *y_i_* of **Y** are the fitness values corresponding to the *i*th row of **X**, our goal is to estimate the parameters *β*_0_, *β_i_, β_i,j_, β_i,j,k_* of the fitness landscape model in Equation (2), such that for any row *x_i_* of **X**, *f*(*x_i_*) ≊ *y_i_*.

To do this we construct a matrix **X**\* ∈ {0,1}^*n*×*p_int_*^, where 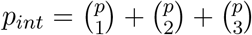, containing a column for each gene, pair of genes, and triplet of genes. Specifically, to construct **X**\* we extend **X** by adding the following columns. For every pair of columns *i, j* and triplet *i,j,k* we add an interaction columns *X_i,j,k_* by taking the element-wise product of the columns *X_i_* and

Given the interaction matrix **X**\* ∈ {0,1}^*n*×*p_int_*^ we estimate effects by solving the square-root lasso, as defined in [57], by minimising the difference between the predicted and actual fitness values, subject to regularisation.

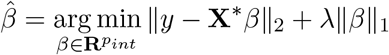

### 2.2. Cyclic Linear Regression

Our approach to lasso regression is based on a cyclic coordinate descent algorithm from [21], as described in [61]. This method begins with *β_j_* = 0 for all *j* and updates the beta values sequentially, with each update attempting to minimize the current total error. Here this total error is the difference between the effects we have estimated and the fitness we observe. Where *y_i_* is the ith element of **Y**, *β_j_* is the jth element of *β*, and *x_ij_* is the entry in the matrix **X**\* at column *j* of row *i*, the error is the following:

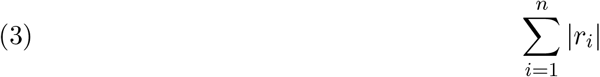

where the residuals, *r_i_*, are the following:

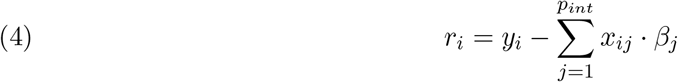

The error affected by a single beta value can then be minimized, using lasso regularisation, by updating *β_k_* as follows:

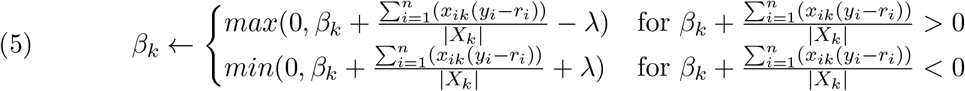

We adjust this to instead solve the square-root lasso ([6]) using Equation (6).

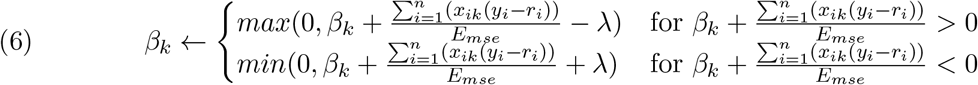

where

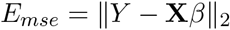

We cyclically update each *β_k_* until the solution converges for a particular λ, reduce the value of λ, and repeat. To reach the genome-scale we avoid unnecessarily considering most interactions (Section 2.4), compress the matrix (Section 2.5), and parallelise the process (Section 2.6).

### 2.3. Choosing Lambda

The lasso penalty requires a regularisation parameter λ, which effectively decides how large an effect has to be before it will be included in the model. This can range from allowing all values (λ = 0) to allowing none (λ → ∞). Choosing the correct value of λ is essential if we want to include only the significant effects and avoid over-fitting noise. For the standard lasso this is typically done by choosing an initial value sufficiently large that all beta values will be zero and gradually reducing λ, fitting the model for each value until a stopping point chosen with *K*-fold cross-validation [20]. Cross-validation requires fitting each λ value *K* times, however, significantly increasing the running time. The square root lasso performs well with an easily calculated λ value, independent of the standard deviation of noise [6]. We use this lower limit of 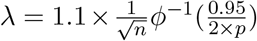, where *ϕ* is the probability density function of the standard normal distribution and *p* is the number of columns of the **X**\* matrix. Note that Belloni, Chernozhukov, and Wang [6] actually use the penalty 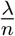, whereas we use λ directly as in Tian, Loftus, and Taylor [57]. To reach the same minimum value as in [6], our equation differs from theirs by a factor of *n*.

### 2.4. Pruning

We implement the branch pruning (Equation (7)) and working set (Algorithm 1) algorithms from WHInter [39] to avoid considering unnecessary effects at each value of λ.

The pruning algorithm determines whether any interaction with a particular effect *i* can be non-zero at the current λ, and if not removes all such interactions *i, j*. Effects that may have a sufficiently large interaction are instead included in the working set. We give a brief overview of this algorithm here, and refer the reader to [39] for further details.

For the square-root lasso penalty we keep track of *ω* = ||*r*||_2_ at each iteration, where *r* = *Y* – **X***β*. For a particular column index *x* in **X**\*, we further define:

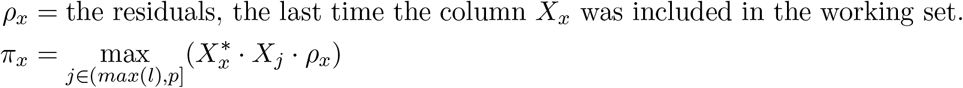

where *l* is either the index of a single column of **X**, or the set {*i,j*} for the interaction column 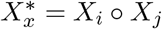. The upper bound *η*(*x*) for any interaction with the column 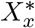 is then:

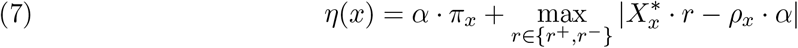

where:

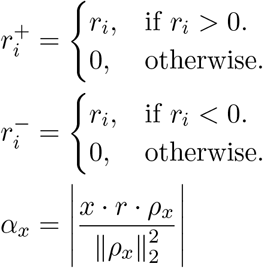

According to the Karush-Kuhn-Tucker conditions for the square-root lasso, no effect can have a value less than |λ · *ω*| [57]. We can therefore ignore any interactions whose effect has an upper bound below |λ · *ω*|. The working set contains the columns that have not been ruled out.

This fast step allows us to rule out most effects without even calculating them. Many interactions even among these columns will still never be updated at the current value of λ however. As in [39] we further reduce the problem by calculating the *active set*, the subset of the working set that will be updated in the current iteration. To do so, we iterate through the entire working set one time, calculating all interactions and updating *ρ* and *π* for each, and adding those that are significant enough to the active set (Algorithms 1 and 3).

#### 2.4.1. Active-set for Pairwise Interactions

For each pair *i,j* of effects that have not been ruled out, we need the sum of their row’s residuals *r* · *X_i_* · *X_j_*. As in [39], rather than taking all the columns and reading through both to find the places they overlap we store the matrix in both column and row major versions and read through only the first column **X**_*i*_. All interactions are found by reading the row *k* for each non-zero entry in *X_i_*. Since the matrix is stored in a sparse format, we find all pairwise interactions with **X**_*i*_ in 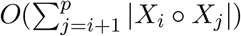 operations. This is further improved in our implementation by calculating a reduced row-major version of *X, X*′, containing only the effects present in the working set. Once the active set has been calculated, we solve the regression problem for the current λ with Algorithm 2. In `Pint` we solve the main effects *β_i_* alone first, followed by pairwise effects *β_i,j_* (and finally three-way effects *β_i,j,k_*). This ensures pairwise effects are only used to explain variance that cannot be fit using main effects. Similarly, three-way effects should only be introduced where pairwise effects are inadequate.

##### Algorithm 1: Identify Active Set (WHInter version, pairwise interactions only).

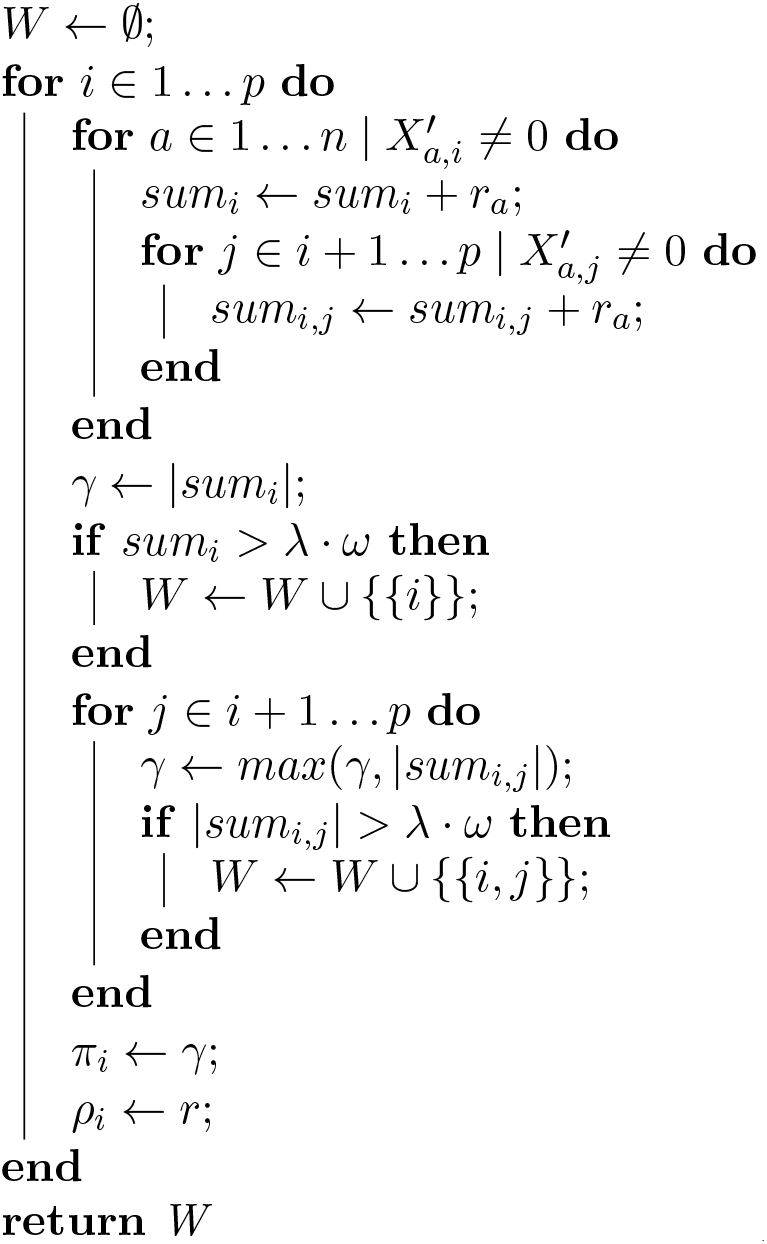

##### Algorithm 2: Sequential Cyclic Sub-problem Algorithm.

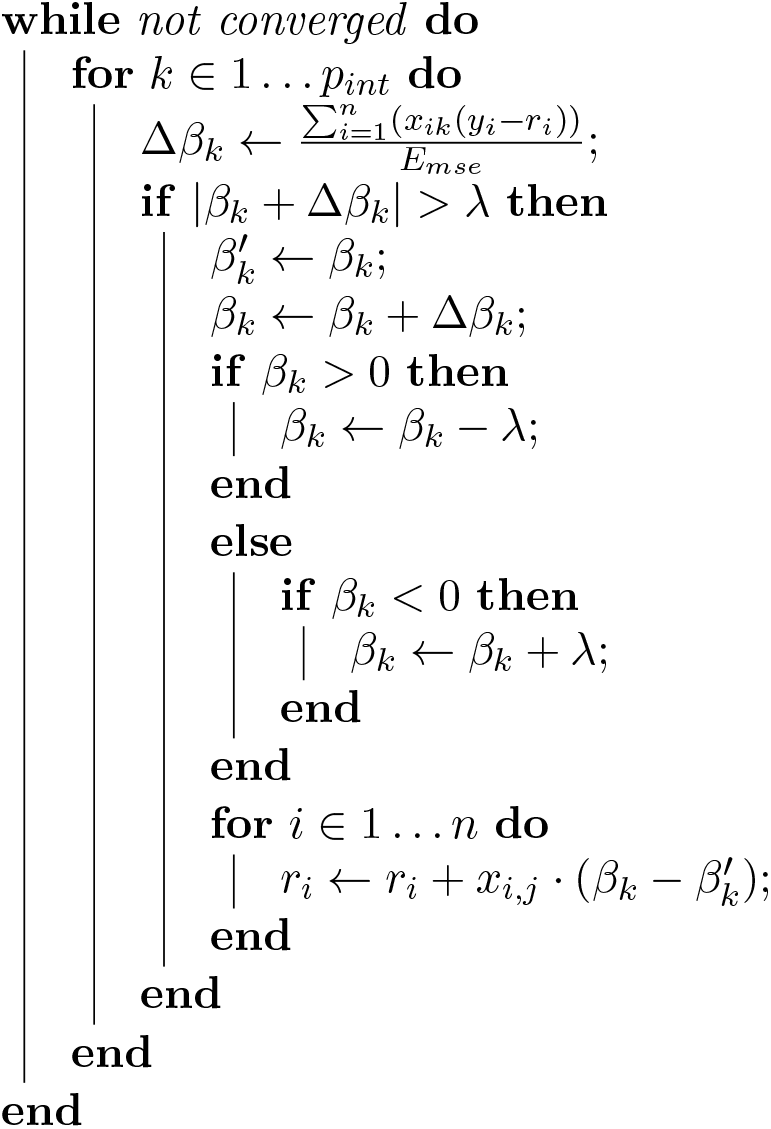

#### 2.4.2. Active-set for Three-way Interactions

WHInter’s pruning algorithm (Algorithm 1) can be extended to three way interactions with a second-level pruning step while updating the active set (Algorithm 3). The three-way active set *W* can then be solved as before using Algorithm 2. Since *η*(*x*) requires the column 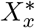, to be calculated, and we are now using interaction columns *x* = {*i,j*}, we keep a cache of every interaction 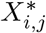 calculated so far and re-use them. For sufficiently small λ this may become the majority of **X**\*, so we compress these columns using Simple-8b (Section 2.5). The upper bound *η*({*i,j*}) may be re-used and should also be cached.

##### Algorithm 3: Identify Active Set (Pint version, three-way interactions)

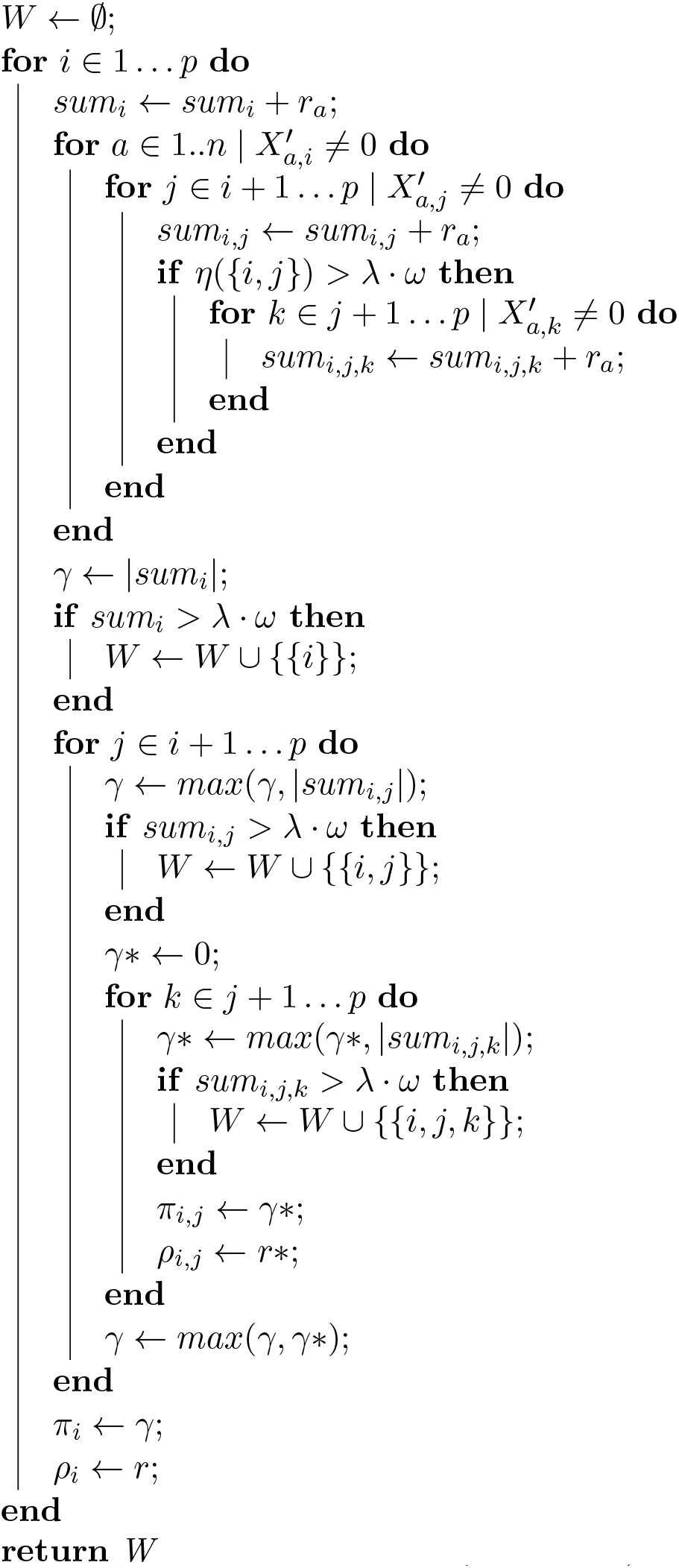

### 2.5. Compression

In our method (Algorithm 3), the input matrix **X** is accessed frequently, iterating through both rows and columns. Because the matrix is not prohibitively large, we store it as both a column and row-major *uncompressed* sparse matrix. The active set, on the other hand, can be as large as **X**\*. We considerably increase the number of possible non-zero effects by storing this only as a set of Simple-8b compressed columns (Figure 2 (c)). Because we read the columns sequentially, we replace each entry with the offset from the previous entry. This reduces the average entry to a relatively small number, rather than the mean of the entire column. These small integers can then be efficiently compressed with any of a range of integer compression techniques (Figure 2), a subject that has been heavily developed for Information Retrieval. We compare a number of such methods, including the Simple-8b algorithm from [53] (which we implement and use in our package) in Appendix A.

**FIGURE 1.**
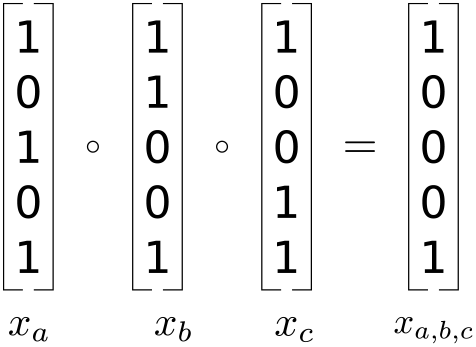
Creation of three-way interaction effect columns

**FIGURE 2.**
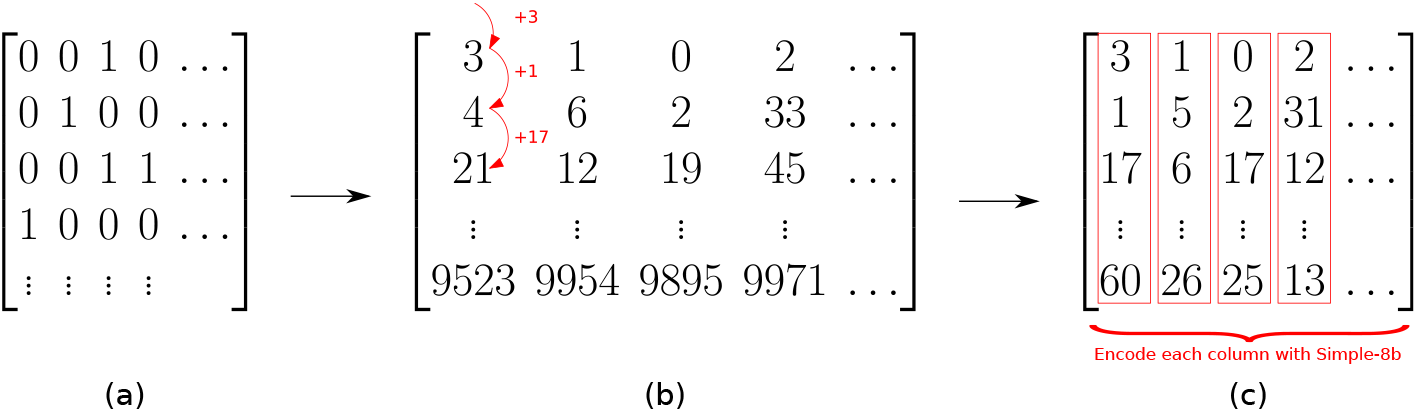
Matrix Compression. Given a full matrix (a), we reduce it to the indices of non-zero entries (b), then the compressed difference between these (c). Arrows represent transitions between different representations.

### 2.6. Parallelisation

The three components of the algorithm, pruning, active set calculation, and solving the sub-problem, can all be done in parallel. Pruning and active-set calculation are trivially parallelisable, and performance scales well as long as each thread is given a large separate chunk of work. In practice this means dividing the matrix into several continuous chunks for the pruning step, one for each thread. For the active set calculation we calculate all two and three way interactions with a particular column on the same thread. As well as keeping each threads workload sufficiently large, this means cached interaction columns and upper bounds can be kept thread-local.

The sub-problem (Algorithm 2) can also be parallelised, and performs well when the active set is sufficiently large and shuffled before each iteration [11]. Parallelising updates to a small active set can significantly harm performance however. In practice this presents a number of difficulties. The active set contains only the columns that would be non-zero at the current value of λ, and this is initially high, with the active set containing very few elements. It is therefore not worth parallelising until λ reaches a value where sufficiently many non-zero effects are allowed. In our testing we often reached the final value, 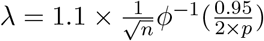, before this occurred.

Furthermore, as λ decreases calculating the active set quickly dominates the running time. While not parallelising the sub-problem calculation theoretically limits the best-case performance of our method, it is a minor limitation in practice. We therefore keep this component single-threaded in our implementation in `Pint`.

We demonstrate the parallel scalability using a simulated data set of *n* = 8,000 rows and *p* = 4, 000 columns, running `Pint` until the first 500 pairwise or main effects have been found and recording the median running-time out of five runs. As we see in Figure 3, performance scales up to an 8 × speed-up using 32 threads on 16 cores across two CPUs, which is typical for a highly parallel task running on CPU(s) with shared memory [32].

**FIGURE 3.**
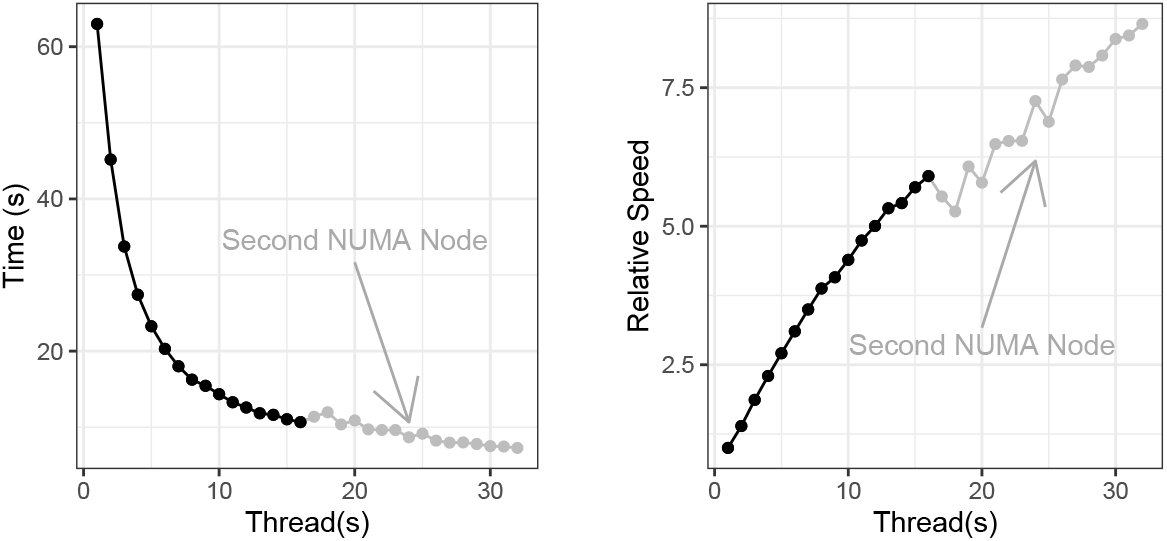
Running time on an increasing number of threads. Note that performance initially decreases with only a small number of threads on a second NUMA node. Tests were performed using two Intel Xeon Gold 6244 CPUs

### 2.7. Approximate Hierarchy

We include an approximate method to enforce a strong hierarchy by only allowing interactions between positions that have at some point had non-zero main effects. This is done by ignoring all interactions *i, j*, where one of *i* or *j* has a main effect strength of zero. In practice, this amounts to replacing the pruning step in *Section* 2.4 with one that simply includes main effects the first time they are assigned a non-zero value.

Doing so significantly reduces both the running time and memory use. We demonstrate this with a test case from the simulated data set, where *n* = 4, 000, *p* = 2, 000, containing 500 pairwise effects, each of which is composed of two main effects. Running to the lower limit for *λ* without the approximate hierarchy constraint takes 42.5 seconds using eight SMT threads across four cores, with peak memory use of 6.87 GB. Adding the approximate hierarchy constraint, running time reduces to 9.98 seconds, and peak memory use to 2.32 GB. Since the running time and memory use depend on the number of main effects rather than *p*, these differences will only increase with larger values of *p*.

To measure the effect this has on accuracy, we run until 100 effects have been found on simulated data sets of *n* = 4, 000, *p* = 8, 000. These contain 40 main effects and 200 pairwise effects (see Section 2.10.1 for details). Note that no effort is made to enforce a hierarchy in the simulation, the components of pairwise effects are chosen randomly. Among predicted pairwise effects we notice a slight drop in recall, and a smaller increase in precision (Fig. 4). This drop in recall may be acceptable in cases where the running time or memory use of the unconstrained method are prohibitive.

**FIGURE 4.**
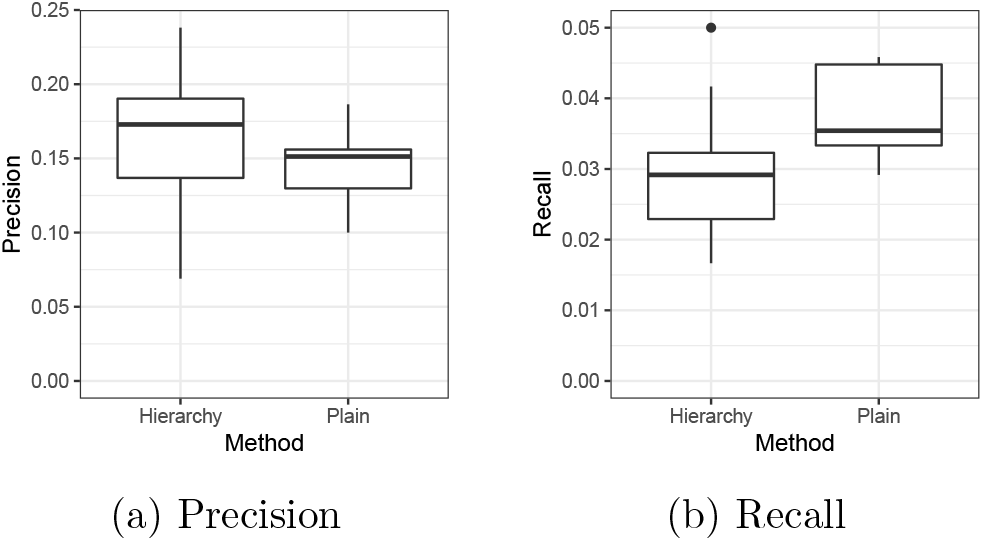
(a) Precision, and (b) recall with and without the approximate hierarchy constraint. ‘Hierarchy’ is with, ‘Plain’ is without. Only pairwise predicted effects are included.

### 2.8. Identifying Identical Columns

We include an option to ignore identical columns in the interaction matrix **X**\*. These may be either direct columns of the input matrix **X** or interaction columns. We do so by computing the 128-bit hash of the column’s non-zero entry positions using XXHASH [12]. All newly-considered columns have their hashes compared to those of previous columns, duplicates are placed on a list of known-duplicate columns and never included in the active set. This avoids spreading out an effect across multiple identical columns. Note that for non-binary matrices columns will be considered identical when the indices of their non-zero entries are the same, even if these entries differ.

### 2.9. Non-binary Matrices

Real values may optionally be included in the matrix **X** ∈ ℝ^n×p^, rather than strictly binary **X** ∈ {0,1}^*n*×*p*^, at the expense of running time. When real-value inputs are used, we maintain a vector *V_k_* of the values for each column of the input matrix **X**_*k*_. In working set calculations, we substitute *v_j_x_i,j_*, or *v_j_x_i,l_* where *l* is an interaction between *j* and *k.* In pruning we avoid actually calculating these values, instead we consider an upper bound on the possible interactions. We store the largest value in each column *k* as 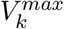, and the largest value overall as 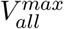. The largest possible interaction with column *k* is then:

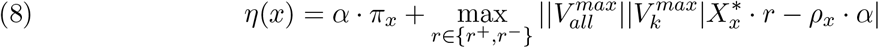

for pairwise interactions, substituting 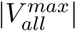 for 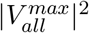 in three-way interactions.

For large values of 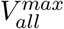 or 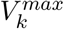 this may include considerably more effects in the active set than if **X** were binary. In both the active-set calculation and the final regression step, we use the real value *v_j_x_i,j_* in place of the binary value *x_i,j_*.

### 2.10. Data

We prepared one simulated and two experimental data sets to evaluate our method and test the scalability of our implementation. The first is the InfectX siRNA perturbation screen [54] in which siRNAs are applied to an infected human cell line. We predict off-target effects across the entire exome, and use these for our analysis. The second data set contains single nucleotide variants (SNVs) from 259 isolates of *Pseudomonas aeruginosa*, and associated minimum inhibitory concentration (MIC) of Ciprofloxacin.

#### 2.10.1. Simulated Data

To evaluate the accuracy of our method `Pint`, we use benchmarks similar to Elmes et al. [16]. To begin with, we take simulated a simulated matrix **X** ∈ {0,1}^*n*×*p*^ resembling siRNA off-target predictions for *n* siRNAs across *p* genes. We randomly assign effects to some of the silencing of some individual and pairwise combinations of the *p* genes to produce effects *β_i_*, and *β_i, j_*. Our simulations differ from [16] in that we do nothing to ensure our pairwise effects are composed of existing main effects (i.e. we do not enforce a hierarchy). Taking the cumulative silencing effects ∑_*i*_ *X_i_β_i_* + ∑_*i,j*_ *X_i_X_j_β_i,j_*, we add random noise from a normal distribution to produce a response vector *Y*, ensuring a signal-to-noise ration of 5.

We simulate three data sets, one with *n* = 1,000 siRNAs and *p* = 100 genes, one with *n* = 8,000 siRNAs and *p* = 4, 000 genes, and one with *n* = 1, 000 siRNAs and *p* = 20,000 genes. The first represents an ideal scenario, with 10 siRNAs per gene at an easily tractable scale. The second is the largest set we are able to run with `glinternet`, and has only two siRNAs per gene. The third represents the worst case, where *p* ≫ *n*. Each simulation in the *p* = 100 set contains 10 main effects and 50 pairwise effects. The *p* = 4,000 simulations contain 40 main effects and 200 pairwise effects. The wide *p* = 20, 000 simulations contain 100 main and 500 pairwise effects.

We simulate a additional sets containing three-way interactions *β_i,j,k_*, such that the cumulative effect is 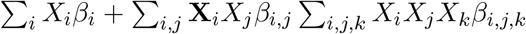 These sets are created with 10 main effects, 20 pairwise effects, and 20 three-way effects. We create sets where *p* = 100, and *p* = 200, with signal to noise ratios of 2, 4, and 8. In each case *n* = 10 × *p*.

We attempt to learn the gene silencing effects from the off-target matrix **X** and the response *Y*.

#### 2.10.2. InfectX siRNA Data

To demonstrate our method on real genome-scale data, we use the mock group from InfectX [46]. This set contains 6, 703 siRNA perturbations (excluding control wells and pooled siRNAs). Off-target effects are predicted using RIsearch2 [2], which includes a gene whenever there is a match between the siRNA seed region and some component of an mRNA for that gene (taken from [26]). We use an energy cut-off of −20 and match the entire siRNA (rather than only the 3’ UTR) as suggested in [2].

We then form the 6, 703 × 19,353 matrix of off-target effects with columns for each gene, and rows for each siRNA as in [16]. An entry *i, j* in this matrix is one if and only the predicted effect of siRNA *i* on gene *j* is greater than zero. All other entries are zero. Our fitness vector **Y** is the result of B-scoring then Z-scoring the number of cells in the well, to remove systematic within-plate effects and experimentally introduced cross-plate biases. B-scoring corrects for biases across the entire plate, Z-scoring then normalises each well’s score with respect to the rest of it’s plate.

#### 2.10.3. Antibacterial Resistance

SNVs from 259 isolates of *Pseudomonas aeruginosa* were sequenced using Illumina technologies (IPCD isolates on MiSeq and QIMR isolates on HISeq). SNVs from raw reads were mapped to the reference genome PAO1 using Bowtie2 (v. 2.3.4) [31] read aligners. Variant reports were then sorted into a table, set up so that each isolate was represented as a row and the presence / absence of each SNV was along the columns. Only genomes that had associated MIC values were included. We removed SNVs that occur less than five times, resulting in a table of 259 rows and 174, 334 columns.

*P. aeruginosa* genome sequences were selected from strains for which MIC values (Ciprofloxacin) were known. 167 genomes were sourced from the publicly available IPCD International Pseudomonas Consortium Database [27] and 92 genomes were from QIMR Brisbane Australia [30]. The IPCD data consisted of 2 x 300 bp MiSeq reads whilst the QIMR data was 2 x 150 bp reads. The MIC values were obtained as a combination of e-test strips [47] and plate-based assays [48, 49].

## 3. Results

In this section, we summarize the results of a simulation study we carried out to compare our method against existing approaches. We also demonstrate our method on two large-scale experimental data sets. We include these as reasonable examples of cases in which our method is applicable and validate the results by comparing them with known effects in the String and NCBI Gene databases [56, 8].

### 3.1. Simulation Performance

Our method aims to have comparable precision and recall to the best performing approach known to us [16] while scaling to much larger data sets. we compare precision and recall with `glinternet`, the most accurate of the methods tested in Elmes et al. [16], and WHInter, a recent fast method based on the idea of limited working sets [39]. We use the simulated data described in Section 2.10.1, and consider only whether a method is able to correctly identify which effects are present, not whether the predicted effect strength is correct.

All methods use regularised regression in some form or another, and for a fair comparison we use the same parameters wherever possible. In each case we instruct the method to stop after the number of effects found matches the number we simulated, 60 in the small sets, 240 in the large sets, and 600 in the wide sets. We assume there is no bias β_0_, and use a convergence threshold of 1%. Although we give the same parameters, all methods return more non-zero effects than we request, to varying extents. In particular, `glinternet` typically gives nearly double the requested number. To keep the running time manageable we do not use crossvalidation in `glinternet`. `Pint` and `glinternet` were run in parallel everywhere except the smallest benchmark, using 16 SMT threads across 8 cores on one CPU. Note that a significant portion of `glinternet`’s running time is single-threaded, and further increasing the number of threads has no noticeable impact on performance. The smallest set is not large enough to benefit from parallelisation in `Pint`, and is run on a single thread. WHInter’s implementation is single-threaded. We further compare the running time of each method. Three-way effects are excluded since neither `glinternet` nor WHInter are able to produce them.

In the *n* > *p* tests we find `Pint` and WHInter perform comparably in terms of precision and recall (Figures 5a, 5b, 5d and 5e). This is unsurprising considering both are solving similar regression problems with the same parameters. `glinternet` typically has lower precision and higher recall than the others, which we would expect since it returns significantly more effects than requested.

**FIGURE 5.**
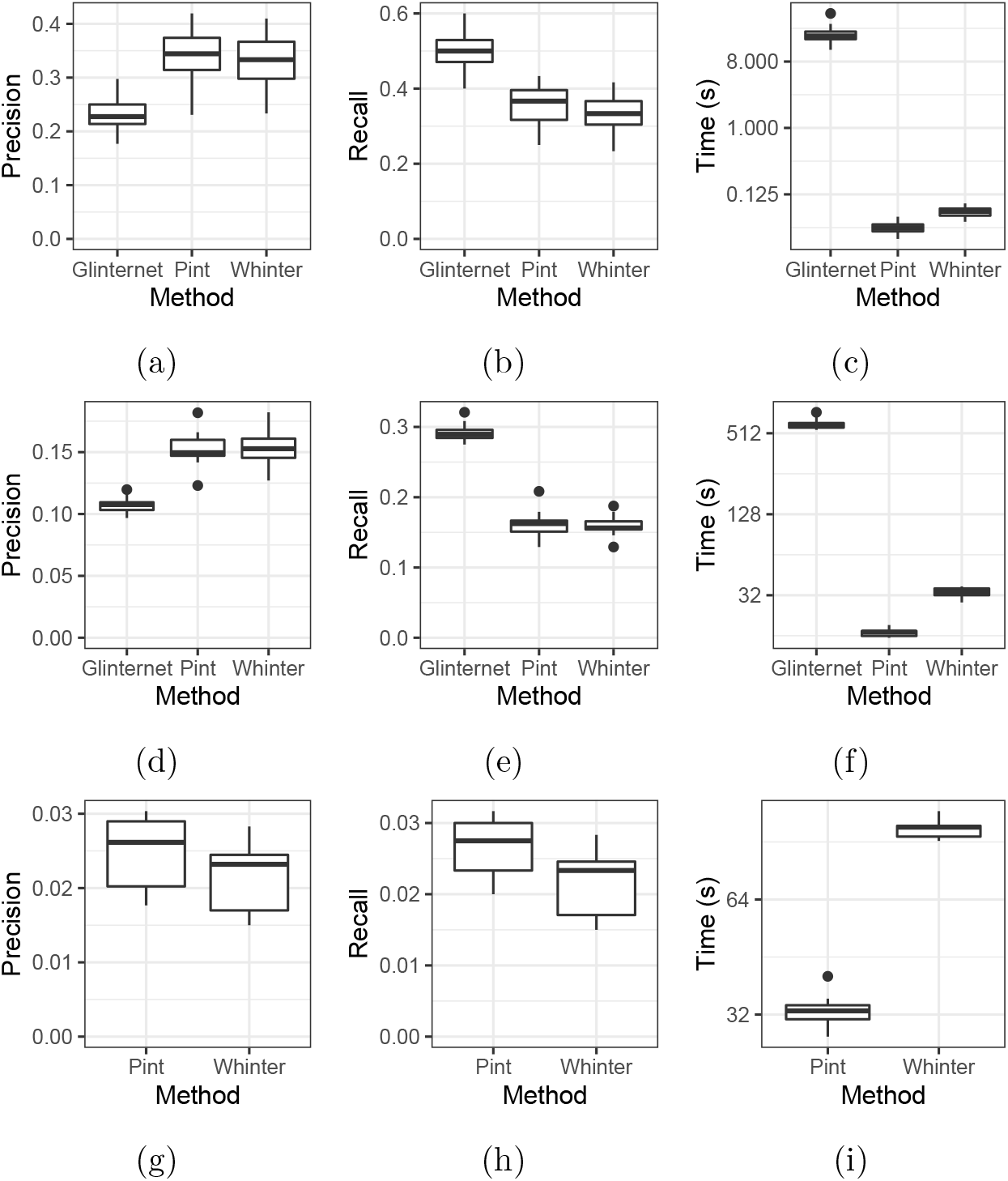
Searching for interactions with `glinternet` vs. WHInter vs. `Pint`. Figures 5a to 5c *p* = 100, *n* = 1, 000. Figures 5d to 5f *p* = 4,000, *n* = 8,000. Figures 5g to 5i *p* = 20, 000, *n* = 1, 000. Running time is measured using one thread (c) or 16 (f,i) on one Intel Xeon Gold 6244 CPU, and shown on a log scale.

In all test cases, we see an improvement in running time compared to WHInter, the fastest method currently available. Using the larger data sets (Figures 5f and 5i) this is because our method runs in parallel, whereas WHInter does not, yielding a significant performance improvement. In the small test cases (Figure 5c) the majority of the used memory fits into the cache on a single core, and we do not see a large improvement from parallelisation. We nonetheless find that `Pint` is faster than WHInter, perhaps due to less efficient cache usage in the latter.

In the wide dataset benchmarks (Figures 5g to 5i) `Pint` has marginally higher precision and recall than WHInter. Since we do not enable the identical column detection described in Section 2.8, this can only be because `Pint` uses the square-root lasso penalty, whereas WHInter uses the classic lasso. `glinternet` takes prohibitively long and is excluded. While the overall precision and recall are particularly poor in this test, with precision and recall below 3%, it is worth noting that the strongest predicted results are more accurate. Considering only the 10 strongest predicted effects in each test, 38 out of 100 effects are correct.

#### 3.1.1. Effect Strength

Effect strength is a strong predictor of accuracy. We quantify this using the simulated three-way effect datasets from Section 2.10.1, finding the first 100 effects in all cases. Sorting predicted effects by their proposed strength, we find that the strongest effects are overwhelmingly true positives. Using absolute effect strength |*β_i_*| as the predictor and plotting the receiver operating characteristic curve, we achieve an area under the curve of 0.93 (Fig. 6). Similar results are reproduced with main, pairwise, and three-way effects separately, which achieve AUC of 0.95, 0.92, and 0.93 respectively. A predicted effect strength threshold can therefore be used to decide the trade-off between precision and recall. Moreover, since strong effects are exactly those that are allowed at a large value of λ, the non-zero effects found in the earliest iterations are the most likely to be correct. In cases where low recall is permissible, limiting the number of non-zero effects (and therefore halting the method at a high λ) not only reduces running time but produces a larger fraction of true positives.

### 3.2. Three-way Effects

In cases where three-way effects are present, we find that including them in the search improves not only the overall fit, but the accuracy of pairwise predictions. This makes sense when we consider that the alternative is to attempt to fit the signal produced by three-way effects to closely matching pairwise effects, which are unlikely to be correct.

To quantify this, we consider the three-way simulations from Section 2.10.1. we run `Pint` until 100 effects are found on all data sets. Precision and recall were near zero for three-way effects, since only one out of 99 was a true positive. This results in a slight drop in total precision and recall when three-way effects are included (Figs. 7c and 7d). There were, however, 11 predicted three-way effects that were indistinguishable from a true positive. That is to say that the *i, j, k* interaction column *X_i_* · *X_j_* · *X_k_* is identical to some other column 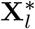, where l is a true effect.

As noted in Section 3.1.1, the small predicted effects are the most unreliable. We therefore focus on the most promising candidates by considering only those with a predicted strength greater than the standard deviation of predicted effects within the same set, *β_i_* > *sd*({*β_k_*|*β_k_* ≠ 0}). Across all ten datasets, the 39 predicted large main effects were the same in both the pairwise-only and three-way cases, and all were correct. There were 27 common large pairwise effects, all correct. The pairwise-only case proposes five more large pairwise effects, of which only one is correct, whereas the three-way case proposes four large three-way effects. While none of these are correct, they are all indistinguishable from some true effect.

Among large effects, the inclusion of these three-way effects reduced the number of large false positive pairwise effects proposed (Fig. 7a) with a negligible effect on recall (Fig. 7b).

### 3.3. InfectX siRNA Data

We run our lasso model on the InfectX data (Section 2.10.2) allowing all pairwise and three-way interactions and halting at the end of the iteration once 100 non-zero effects are found. Only the combinations with non-zero predicted effects are then included in the matrix **Z**. We then fit the fitness values **Y** to this matrix using least-squares regression **Y** ~ **Z***β*, using these unbiased estimates and p-values as our final result. The resulting fit has an adjusted R-squared of 0.13 and an AIC value of 18,184.82 We summarise the five most significant (according the p-value of the fit **Y** ~ **Z**) estimated effects in Table 1.

**TABLE 1.**
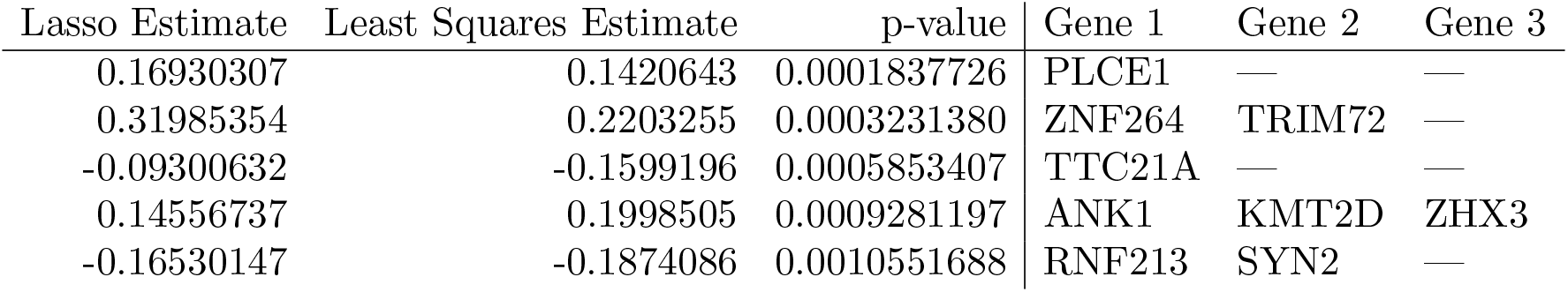
InfectX most significant proposed effects.

Among these effects, the three-way suppression of ANK1, KMT2D, and ZHX3 is particularly plausible. KMT2D is a known tumor suppressor, and mutations are common in lymphoma [63, 41]. ZHX3 is a transcriptional repressor, and in particular a failure of ZHX3 expression may be a cause of hepatocellular carcinoma [62]. Changes and failure to express KMT2D and ZHX3 respectively are associated with cancer development, and a significant increase in cell growth after suppressing both is consistent with these functions. ANK1 attaches integral membrane proteins to the cytoskeleton [59], and what role, if any, it plays is unclear.

The pairwise effect suppressing TRIM72 and ZNF264 could plausibly affect cell survival, as could suppressing PLCE1. PLCE1 is believed to play a role in cell survival and growth, and its suppression could have a significant effect on its own. It is unclear, however, how its suppression could have a positive effect on cell count [59]. TRIM72 plays a central role in cell membrane repair, and its suppression could easily affect fitness. ZNF264 may be involved in transcriptional regulation, and may have an interacting effect, although there are no known interactions between ZNF264 and TRIM72 [59].

The remaining two effects are TTC21A on its own, and RNF213 combined with SYN2. These genes are known to be involved in sperm function, vascular development, and neurotransmitter regulation respectively [36, 59]. We are not aware of any way in which RNF213 and SYN2 interact, or how either of these effects might affect cell growth or survival.

### 3.4. Antibacterial Resistance

As explained in Section 2.10.3, our antibacterial data is pre-processed to remove variants present in less than five cases. The remaining 174, 334 columns are included in the model. We then fit the model *Y* = *X*_2_*β* + *ϵ*, where *Y* is the MIC *log*_2_ phenotype (indicative of Ciprofloxacin resistance). We include all pairwise interactions in *X*_2_, and stop after 50 non-zero effects are found. These effects are included even with a large regularisation value λ, and are the most likely to be true positives (see Section 3.1.1). Again creating a *Z* matrix with only the non-zero columns fitting *Y* ~ *Z*, we get a least-squares estimate with an Adjusted R-squared of 0.23.

The 50 non-zero effects involve 47 variants with 19 repeats and 16 pairwise effects (Appendix, Table 2). Of the pairwise effects, two pairs include the non-synonymous variant change that results in a Leu523Gln change in PA3054. PA3054 encodes a putative carboxypeptidase with the peptidase_M14 domain occurring between bases 24-634. Extracellular degradation of antimicrobials has been associated with increased production of M14 carboxypeptidases [55]. All other pairwise interactions identified involved synonymous variants. The most common of these was an A to G variant in PA3460, codon 537 Leu, found in 50% of the interactions. PA3460 encodes an acetyltransferase that is able to possibly modify Fluoroquinolones, reducing bacterial susceptibility to Ciprofloxacin [51]. The second most common synonymous variant was another A to G change in PA3709 encoding Ala 340 of a probable major facilitator superfamily (MFS) transporter protein. Overexpression of efflux pumps that include MFS transporters are associated with increased resistance to antibiotics [15, 40].

There were 34 variants that were characterised as contributing to Ciprofloxacin resistance. Of these, 16 were non-synonymous changes to proteins that are involved in fluoroquinolone modification, membrane transport or oxidative stress responses. The majority of non-synonymous variants occurred in oxidative stress response genes. An increase in reactive oxygen species (ROS) in response to Ciprofloxacin is well characterised in bacteria [24, 28, 1]. PA5401 encodes an electron transfer flavoprotein (EFT) domain-containing protein. The variant results in an Arg36Cys change in the *β*-subunit of an electron transfer protein whose gene is part of the dgc operon that is involved in choline metabolism and associated pathogenesis [19]. In eukaryotes, EFT is known to produce significant amounts of ROS in the presence of its partner enzyme medium-chain acyl-CoA dehydrogenase (MCAD) [52]. Therefore, non-synonymous variants in PA5401 could result in changes to pathogenesis or ROS amounts. Our method also identifies effects for PA0117, pauD2, and gloA1, all of which are involved in glutathione production. Glutamine is a precursor of glutathione; glutamine and ascorbic acid have been found to provide substantial protection against Ciprofloxacin susceptibility in *Escherichia coli* [24].

Two genes with non-synonymous variants that were identified are involved in membrane integrity. The first is PA3173 with a His93Arg variation that encodes a short-chain dehydrogenase and acts on ubiG and ubiE involved in ubiquinone biosynthesis [29]. ROS accumulation affects membrane systems due to lipid peroxidation [18]. Ubiquinone is lipid-soluble and, therefore, is able to act as a mobile redox carrier within the cellular membrane [22]. Increased production of ubiquinone would reduce membrane damage caused by ROS. The second gene, MviN, with Leu316Met, is involved in peptidoglycan biosynthesis. Fosfomycin is frequently co-prescribed with Ciprofloxacin due to the synergistic activity of the two drugs [60]. However, increased peptidoglycan biosynthesis and cell wall recycling lead to antibiotic resistance [9]. Therefore the Leu316Met variant in MviN could be linked to increased resistance to combination therapy of Ciprofloxacin and Fosfomycin.

Overexpression of efflux pumps is a known contributor to increases in MICs for *P. aeruginosa*. Identified was a variant Lys329Gln in mexX that encodes a resistance-Nodulation-Cell Division (RND) multidrug efflux membrane fusion protein MexX precursor.

In total, 13 of the 16 non-synonymous variants have possible contributions to Ciprofloxacin resistance.

## 4. Discussion

Genotype-phenotype data sets have recently become available at a never before seen scale. In principle, it is possible to infer not only the effect of individual genomic variants within such data, but of pairwise and higher order combinations of their effects. While this has been shown to work in theory, and a number of tools have been developed that work on a smaller scale, there is a shortage of effective methods for human genome-scale data, and no method we are aware of includes three-way interactions. In this paper we present a lasso regression based method for such large-scale inference of pairwise and three-way effects.

Our method effectively performs coordinate descent square-root lasso regression on a matrix containing all pairwise and three-way combinations of the input data. We expand upon the method used in [39], with a number of improvements. We update the working set in parallel, resulting in a significant speed improvement. We extend the method from two-way interactions to three-way, adding pruning of pairwise effects. The active set is compressed with simple-8b, significantly reducing memory use and improving the running time with a large number of nonzero effects. We extend the method to include non-binary inputs in the **X** matrix, introduce an optional approximate hierarchy constraint that can be used to further reduce running time and memory use, and add detection of identical columns. Finally, we solve the square-root lasso instead of the lasso, giving us a well-defined stopping point.

We compared the accuracy and running time of our work to `glinternet`, the best of the methods we used previously [16] and WHInter, the fastest running method we are aware of [39]. Our simulations demonstrate comparable accuracy and recall to existing methods, running approximately three times faster than WHInter and 60 times faster than `glinternet` in the largest tests. Considering only the largest effects, our method is able to achieve precision > 30% even in wide data sets where *p* ≫ *n* (Section 3.1). Moreover, the stronger a predicted effect is the more likely it is to be correct (Fig. 6). We therefore expect focusing on large predicted effects will achieve reasonable precision even on extremely large datasets.

**FIGURE 6.**
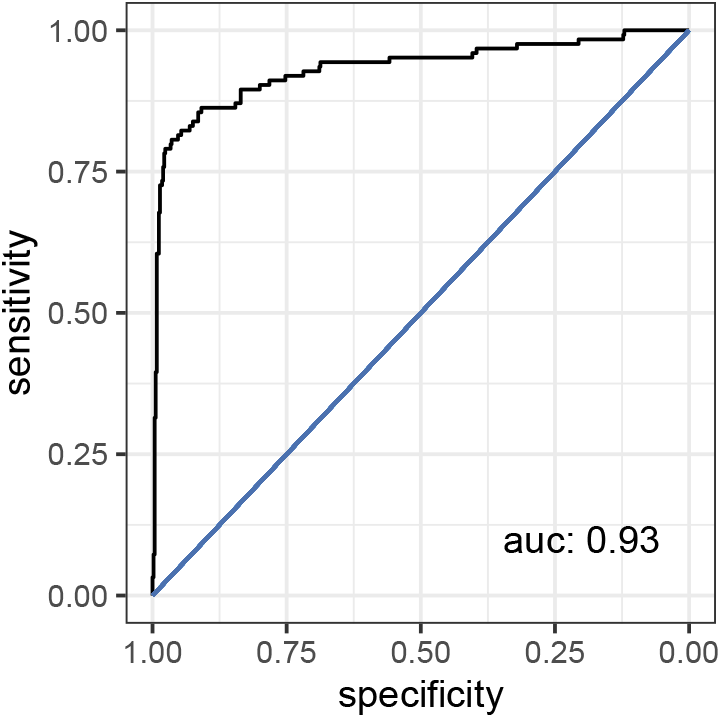
Receiver operating characteristic curve comparing fraction of reported effects that are true positives as the predicted strength varies.

**FIGURE 7.**
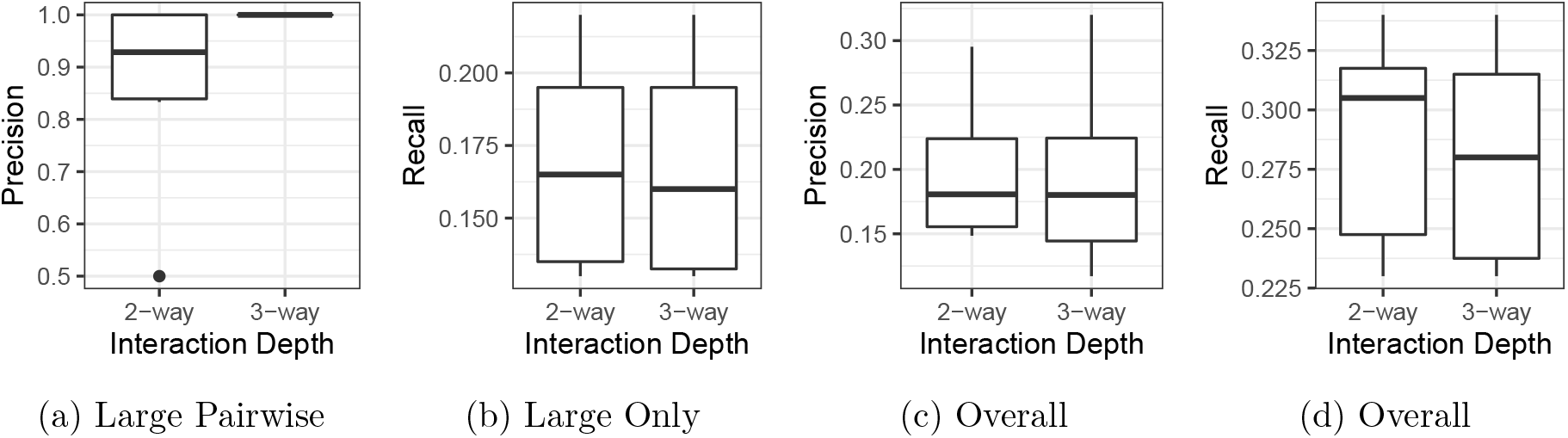
Precision and recall of predicted effects using `Pint`. Data is fit with single, pairwise, and three-way interactions allowed (3-way), or only single and pairwise (2-way). (a) Precision of predicted large pairwise effects. (b) Recall, including main, pairwise, and three-way predictions of predicted large effects. (c, d) Overall precision and recall including small effects.

We also tested our method using two genome-scale real data sets. One is an exome-wide siRNA perturbation screen (*n* ≊ 6, 700 siRNAs and *p* ≊ 19,000 genes). The other measures antibacterial resistance with respect to genetic variations in *Pseudomonas aeruginosa*, and includes over 15 billion possible pairwise interactions. In both cases our method finds a number of plausible interactions.

Despite this success, our method and its implementation in `Pint` have the following limitations. While our method is effective on genome scale data when using only pairwise interactions, running time limits the use of three-way interactions to smaller sets, or only the strongest interactions (running to completion was only possible with *p* ≤ 5, 000 in our testing). Furthermore, the sub-problem given the working set is not solved in parallel. While it is possible to do so, it is actually harmful to performance unless the working set is very large (i.e. many non-zero effects are included).

Note that while we consider only pairwise interactions Section 3.4, it is possible to include three-way effects if we remove columns with less than 30 entries instead. This reduces the input from 174, 334 to 75, 599 columns, and the first 50 interactions can then be found in approx. 80 hours.

The additive interaction model is also an oversimplification of biology. It remains unclear to what extent genetic effects be treated as additive, and ignoring interactions among of more than three items could well be leaving out the most important effects. In this case we may end up spuriously associating phenotype changes with effects that just happen to be present, rather than the true, more complicated, interactions (as demonstrated in Section 3.2). Finally, we cannot distinguish interactions that are present in exactly the same rows of the input matrix. In the siRNA case, if two distinct pairs of genes are simultaneously suppressed by the same siRNAs in all cases, whichever is considered first will likely trump the other.

There are hence a number of opportunities to expand upon this work. While we can avoid including duplicate effects in our model (Section 2.8), we do not detect indistinguishable effects unless they are considered for inclusion in the working set. Moreover, we do not identify effects that are almost, but not quite, identical. Thoroughly accounting for the similarity of effects would further improve the model. Additionally, we chose the square-root lasso penalty partially because it has simpler to compute p-values than the lasso [57]. Implementing this would give unbiased p-values based directly on the lasso results, without requiring a second least squares fit. This is particularly important since the least squares p-values do not account for the column selection process, and are likely to be biased [33].

More generally, we could significantly increase the scale of interaction inference methods by reducing the search space. A more targeted approach estimating distance in 3D space using Hi-C [7] for example, would drastically reduce the time and space requirements, allowing higher order interactions to be considered. While we implement an optional approximate weak hierarchy constrain (Section 2.7), a strong hierarchy would further simplify the problem. It is worth noting however that these are not reasonable assumptions for all applications. Finally, the interactions proposed in Section 3.3 may be worth further investigation.

Our method is implemented in C++, and an R package is provided at github.com/bioDS/pint. Code to reproduce the simulations and benchmarking from Section 3 is provided at github.com/bioDS/lasso_data_processing.

## Appendix A. Compression

We can considerably reduce the size of the active set by compressing the columns. Since we have a sequence of increasing integers we can store only the offset from the previous entry, keeping the entries small. The resulting sequence of (mostly) small numbers can then be effi-ciently stored using integer compression methods. We describe the compression method we use in Appendix A.1 and compare it to other methods in Appendix A.2.

### A.1. Simple-8b

Simple-8b is a non-SIMD compression scheme, with performance comparable to other state of the art methods [53, 38, 58]. While SIMD-based compression schemes can often offer significantly improved compression and decompression speed [34] [53], their implementation is architecture dependant. Simple-8b only requires a CPU be able to efficiently handle 64-bit arithmetic, and does not significantly underperform compared to state-of-the-art SIMD techniques in our testing (Appendix A.2).

Simple-8b is a 64-bit variation of the Simple-9 encoding scheme [3], and stores a sequence of integers in a single 64-bit word. The number of integers stored depends on the size of the largest one, and is indicated by a four bit ‘selector’. The remaining 60 bits are divided into integers of size 1, 2, 3, 4, 5, 6, 7, 8, 10, 12, 15, 20, 30, or 60, with between 240 (only possible if all values are zero) and one integer stored. As seen in Figure 8, this considerably reduces the size of **X**_2_ in our test data (two sets from [16], one with *p* = 100, *n* = 1,000, another with *p* = 1, 000, *n* = 10, 000). In the larger *p* = 1, 000 set, total memory use is reduced by over 85% compared to storing integers directly. It is worth noting that this compression works well even for non-sparse sections of the matrix, since the offsets are extremely small. In an extreme case, we can store up to 240 sequential 1’s in a single 64-bit word.

**FIGURE 8.**
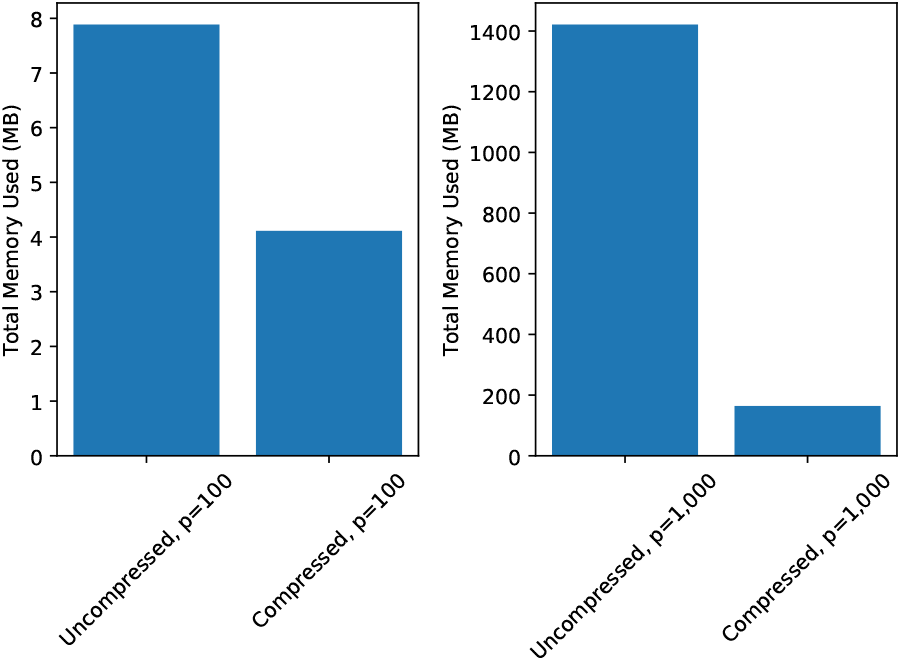
Compression effect on memory use. Note that this is the total peak memory use of the program, not solely the memory used by the matrix **X**_2_. In both cases *n* = 10 · *p*.

### A.2. Comparing Methods

While Simple-8b allows our implementation to be used on any 64-bit CPU, we could also take advantage of SIMD-based methods where such CPU instructions are available. To determine whether this is a worthwhile improvement, we compare our Simple-8b implementation to a number of state of the art alternatives.

Recent work suggests TurboPFor [45] has a particularly high compression ratio [58]. We therefore compare the best performing methods from TurboPFor against our implementation of Simple-8b (Figure 9). The tests are performed using an eight-core (16 SMT threads) Intel Xeon Gold 6244 CPU. To compare these methods, we perform 50 regression iterations on a test data set of *p* = 1,000 genes and *n* = 10, 000 siRNAs. We examine the total time taken for the process, as well as the total memory used and time for the regression function alone (excluding calculating and compressing the interaction matrix).

**FIGURE 9.**
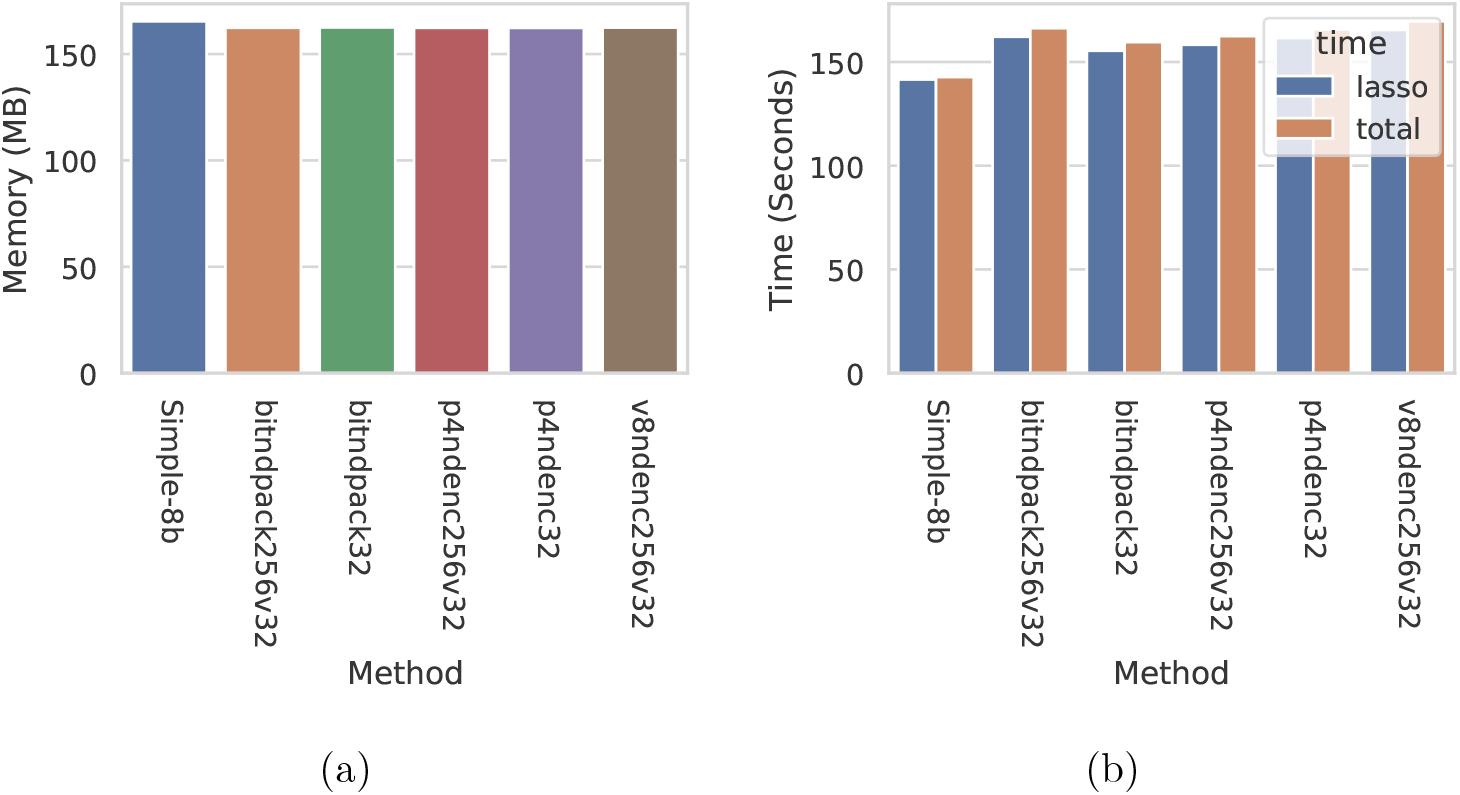
Comparison of Compression Methods. (a) Total memory used, compressing the sparse **X**_2_ matrix with each method. (b) Total time taken and time taken (including compressing **X**_2_) and time taken for lasso regression alone, using each method.

We see that both the time to produce the compressed matrix (seen in Figure 9 as the difference between total time and lasso-only time), and the running time are comparable for all TurboPFor methods.^2^ While every TurboPFor method we tested improved the compression ratio compared to Simple-8b (Figure 9a), we consistently found that the running time was longer (Figure 9b). It is possible that this is a result of the way the columns are being read in each method. Using TurboPFor, we compress and decompress entire columns at a time. With our Simple-8b implementation, we process each 64-bit word separately. This allows us to use the column as it is being decompressed. Avoiding re-reading the column after decompression also allows the entries to be evicted from the cache earlier.

While it is also possible to process compressed words as they are read using the tested TurboPFor methods, there does not appear to be a significant difference in compression that would justify doing so.

## Appendix B. Antibiotic Results

**TABLE 2.** Predicted top 50 SNV effects.

| Lasso Estimate | Least Squares Estimate | pval | SNV 1 | SNV 2 |
| --- | --- | --- | --- | --- |
| -0.024 | -1.76 | 0.01 | 813809 sub. T C | — |
| -0.013 | -5.02 | 0.02 | 3789873 sub. T C | — |
| -0.039 | -1.68 | 0.02 | 4617770 sub. C T | — |
| -0.007 | 9.71 | 0.03 | 5111434 sub. A G | — |
| -0.006 | -2.83 | 0.03 | 4153211 sub. A G | 5719820 sub. A G |
| 0.000 | -1.15 | 0.05 | 6081497 sub. T C | — |
| -0.007 | -2.25 | 0.06 | 4271404 sub. T C | — |
| -0.023 | -2.25 | 0.06 | 63407 sub. T C | — |
| -0.063 | -2.25 | 0.06 | 184347 sub. T C | — |
| -0.001 | -12.62 | 0.07 | 137771 sub. A G | — |
| -0.008 | -2.76 | 0.10 | 3654726 sub. T C | 3869317 sub. A G |
| -0.019 | -1.03 | 0.11 | 1886446 sub. T C | — |
| -0.045 | 13.18 | 0.11 | 442184 sub. C G | — |
| -0.001 | -2.10 | 0.12 | 986363 sub. A G | — |
| -0.003 | -1.84 | 0.13 | 2996864 sub. C T | 3869317 sub. A G |
| -0.030 | -1.60 | 0.16 | 1810265 sub. A C | — |
| -0.012 | 0.76 | 0.20 | 2033361 sub. A C | — |
| -0.009 | -7.03 | 0.22 | 3318136 sub. T C | — |
| -0.038 | 8.19 | 0.23 | 176977 sub. A G | — |
| -0.002 | 2.44 | 0.24 | 172758 sub. T C | — |
| -0.009 | 2.00 | 0.24 | 4003269 sub. T C | — |
| -0.034 | 8.48 | 0.24 | 4153211 sub. A G | — |
| -0.008 | -6.99 | 0.28 | 3419855 sub. T A | 4153211 sub. A G |
| -0.031 | -1.25 | 0.30 | 4847782 sub. A C | — |
| -0.005 | -1.25 | 0.30 | 1383522 sub. A G | — |
| -0.005 | -1.25 | 0.30 | 176977 sub. A G | 5133603 sub. A G |
| -0.004 | 2.36 | 0.30 | 3419855 sub. T A | 4920968 sub. C A |
| -0.040 | 2.44 | 0.31 | 3944454 sub. T A | — |
| -0.004 | -0.84 | 0.40 | 3195104 sub. T C | — |
| -0.028 | -0.42 | 0.40 | 2211528 sub. T G | — |
| -0.013 | 2.14 | 0.51 | 313283 sub. A C | — |
| 0.000 | 0.90 | 0.51 | 3601029 sub. T C | 3869317 sub. A G |
| -0.005 | 4.38 | 0.53 | 3419855 sub. T A | — |
| -0.010 | 2.64 | 0.55 | 5254860 sub. T C | — |
| -0.016 | -0.31 | 0.69 | 2093685 sub. A T | — |
| -0.007 | 1.02 | 0.73 | 3869317 sub. A G | 4687756 sub. A G |
| -0.044 | -0.59 | 0.78 | 4453098 sub. A G | — |
| -0.005 | 0.24 | 0.80 | 3563823 sub. A G | — |
| -0.039 | 1.21 | 0.81 | 4125422 sub. T C | — |
| -0.002 | 0.57 | 0.88 | 3869317 sub. A G | 4966957 sub. C T |
| -0.103 | 0.21 | 0.95 | 3869317 sub. A G | — |
| -0.005 | — | 1.00 | 956627 sub. A G | — |
| -0.001 | — | 1.00 | 5856921 sub. A G | — |
| -0.002 | — | 1.00 | 3869317 sub. A G | 4617770 sub. C T |
| -0.014 | — | 1.00 | 3869317 sub. A G | 4847782 sub. A C |
| -0.008 | — | 1.00 | 4153211 sub. A G | 4453098 sub. A G |
| -0.046 | — | 1.00 | 3869317 sub. A G | 4453098 sub. A G |
| -0.006 | — | 1.00 | 4153211 sub. A G | 5990479 sub. C G |
| -0.009 | — | 1.00 | 3195104 sub. T C | 4153211 sub. A G |
| -0.046 | — | 1.00 | 986363 sub. A G | 3869317 sub. A G |

## Footnotes

1 Fitting interactions in an siRNA screen of 1, 000 genes with ten siRNAs per gene takes several days using ten cores on an Opteron 6276.

2 The compression time is not comparable for all methods. Our Simple-8b implementation compresses columns in parallel, whereas TurboPFor does not. Columns are decompressed in parallel in both cases

## References

[1] Marwa N. Ahmed et al. “Evolution of Antibiotic Resistance in Biofilm and Planktonic Pseudomonas Aeruginosa Populations Exposed to Subinhibitory Levels of Ciprofloxacin”. Antimicrobial Agents and Chemotherapy 62.8 (2018), e00320–18. doi: 10.1128/AAC.00320-18.

[2] Ferhat Alkan et al. “RIsearch2: Suffix Array-Based Large-Scale Prediction of RNA–RNA Interactions and siRNA off-Targets”. Nucleic Acids Research 45.8 (May 2017), e60–e60. doi: 10.1093/nar/gkw1325.

[3] Vo Ngoc Anh and Alistair Moffat. “Inverted Index Compression Using Word-Aligned Binary Codes”. Information Retrieval 8.1 (Jan. 2005), pp. 151–166. doi: 10.1023/B:INRT.0000048490.99518.5c.

[4] Alan Ashworth, Christopher J. Lord, and Jorge S. Reis-Filho. “Genetic Interactions in Cancer Progression and Treatment”. Cell 145.1 (Apr. 2011), pp. 30–38. doi: 10.1016/j.cell.2011.03.020.

[5] N Beerenwinkel, L Pachter, and B Sturmfels. “Epistasis and Shapes of Fitness Landscapes”. Statistica Sinica (2007).

[6] A. Belloni, V. Chernozhukov, and L. Wang. “Square-Root Lasso: Pivotal Recovery of Sparse Signals via Conic Programming”. Biometrika 98.4 (Dec. 2011), pp. 791–806. doi: 10.1093/biomet/asr043.

[7] Jon-Matthew Belton et al. “Hi-C: A Comprehensive Technique to Capture the Conformation of Genomes”. Methods (San Diego, Calif.)58.3 (Nov. 2012), pp. 268–276. doi: 10.1016/j.ymeth.2012.05.001. pmid:22652625.

[8] Bethesda (MD): National Library of Medicine (US), National Center for Biotechnology Information and Citation Key: GeneInternet. Gene [Internet].

[9] Marina Borisova, Jonathan Gisin, and Christoph Mayer. “Blocking Peptidoglycan Recycling in Pseudomonas Aeruginosa Attenuates Intrinsic Resistance to Fosfomycin”. Microbial Drug Resistance 20.3 (June 2014), pp. 231–237. doi: 10.1089/mdr.2014.0036.

[10] João Botelho, Filipa Grosso, and Luísa Peixe. “Antibiotic Resistance in Pseudomonas Aeruginosa - Mechanisms, Epidemiology and Evolution”. Drug Resistance Updates 44 (May 2019), p. 100640. doi: 10.1016/j.drup.2019.07.002.

[11] Joseph K. Bradley et al. “Parallel Coordinate Descent for L1-Regularized Loss Minimization”. arXiv:1105.5379 [cs, math] (May 2011). arXiv:1105.5379 [cs, math].

[12] Yann Collet. xxHash - Extremely Fast Hash Algorithm. July 2022.

[13] Michael Costanzo et al. “The Genetic Landscape of a Cell.” Science (2010).

[14] Kristina Crona et al. “Inferring Genetic Interactions from Comparative Fitness Data”. Elife 6 (Dec. 2017).

[15] Edda De Rossi et al. “The Multidrug Transporters Belonging to Major Facilitator Superfamily (MFS) in Mycobacterium Tuberculosis”. Molecular Medicine 8.11 (Nov. 2002), pp. 714–724. doi: 10.1007/BF03402035.

[16] Kieran Elmes et al. “Learning Epistatic Gene Interactions from Perturbation Screens”. PLOS ONE 16.7 (July 2021), e0254491. doi: 10.1371/journal.pone.0254491.

[17] D. S. (Douglas Scott) Falconer. Introduction to Quantitative Genetics. 4th ed. Harlow, Essex, England: Longman, 1996. isbn: 0-582-24302-5.

[18] Edward E. Farmer and Martin J. Mueller. “ROS-Mediated Lipid Peroxidation and RES-Activated Signaling”. Annual Review of Plant Biology 64.1 (2013), pp. 429–450. doi: 10.1146/annurev-arplant-050312-120132.

[19] Liam F. Fitzsimmons et al. “Small-Molecule Inhibition of Choline Catabolism in Pseudomonas Aeruginosa and Other Aerobic Choline-Catabolizing Bacteria”. Applied and Environmental Microbiology 77.13 (July 2011), pp. 4383–4389. doi: 10.1128/AEM.00504-11.

[20] Jerome Friedman, Trevor Hastie, and Robert Tibshirani. “Regularization Paths for Generalized Linear Models via Coordinate Descent”. Journal of Statistical Software 33.1 (2010). doi: 10.18637/jss.v033.i01.

[21] Wenjiang J. Fu. “Penalized Regressions: The Bridge versus the Lasso”. Journal of Computational and Graphical Statistics 7.3 (Sept. 1998), pp. 397–416. doi: 10.1080/10618600.1998.10474784.

[22] Naoko Fujimoto, Tomoyuki Kosaka, and Mamoru Yam. “Menaquinone as Well as Ubiquinone as a Crucial Component in the Escherichia Coli Respiratory Chain”. Chemical Biology.Ed. by Deniz Ekinci. InTech, Feb. 2012. isbn: 978-953-51-0049-2. doi: 10.5772/35809.

[23] Robert Gaynes, Jonathan R. Edwards, and National Nosocomial Infections Surveillance System. “Overview of Nosocomial Infections Caused by Gram-Negative Bacilli”. Clinical Infectious Diseases 41.6 (Sept. 2005), pp. 848–854. doi: 10.1086/432803.

[24] M. Goswami, S. H. Mangoli, and N. Jawali. “Involvement of Reactive Oxygen Species in the Action of Ciprofloxacin against Escherichia Coli”. Antimicrobial Agents and Chemotherapy 50.3 (Mar. 2006), pp. 949–954. doi: 10.1128/AAC.50.3.949-954.2006.

[25] Alison L Gould et al. “Microbiome Interactions Shape Host Fitness”. Proceedings of the National Academy of Sciences of the United States of America 115.51 (Dec. 2018), E11951–E11960.

[26] GRCh38.P13 - Genome - Assembly - NCBI.

[27] IPCD International Pseudomonas Consortium Database. https://ipcd.ibis.ulaval.ca/.

[28] Peter Ø. Jensen et al. “Formation of Hydroxyl Radicals Contributes to the Bactericidal Activity of Ciprofloxacin against Pseudomonas Aeruginosa Biofilms”. Pathogens and Disease 70.3 (Apr. 2014), pp. 440–443. doi: 10.1111/2049-632X.12120.

[29] Hai-Xia Jiang et al. “Coenzyme Q Biosynthesis in the Biopesticide Shenqinmycin-producing Pseudomonas Aeruginosa Strain M18”. Journal of Industrial Microbiology and Biotechnology 46.7 (July 2019), pp. 1025–1038. doi: 10.1007/s10295-019-02179-1.

[30] Timothy J. Kidd et al. “Pseudomonas Aeruginosa Exhibits Frequent Recombination, but Only a Limited Association between Genotype and Ecological Setting”. PLoS ONE 7.9 (Sept. 2012). Ed. by Sam Paul Brown, e44199. doi: 10.1371/journal.pone.0044199.

[31] Ben Langmead and Steven L. Salzberg. “Fast Gapped-Read Alignment with Bowtie 2”. Nature Methods 9.4 (Apr. 2012), pp. 357–359. doi: 10.1038/nmeth.1923.

[32] Jeff Larkin. OpenMP on GPUs, First Experiences and Best Practices. Nvidia GTC 2018.

[33] Jason D. Lee et al. “Exact Post-Selection Inference, with Application to the Lasso”. The Annals of Statistics 44.3 (June 2016), pp. 907–927. doi: 10.1214/15-AOS1371.

[34] D. Lemire and L. Boytsov. “Decoding Billions of Integers per Second through Vectorization”. Software: Practice and Experience 45.1 (2015), pp. 1–29. doi: 10.1002/spe.2203.

[35] Caitlin Lienkaemper et al. “The Geometry of Partial Fitness Orders and an Efficient Method for Detecting Genetic Interactions”. Journal of Mathematical Biology 77.4 (May 2018), pp. 951–970.

[36] Wangjie Liu et al. “Bi-Allelic Mutations in TTC21A Induce Asthenoteratospermia in Humans and Mice”. American Journal of Human Genetics 104.4 (Apr. 2019), pp. 738–748. doi: 10.1016/j.ajhg.2019.02.020.

[37] Jeffrey B Lyczak, Carolyn L Cannon, and Gerald B Pier. “Establishment of Pseudomonas Aeruginosa Infection: Lessons from a Versatile opportunist1*Address for Correspondence: Channing Laboratory, 181 Longwood Avenue, Boston, MA 02115, USA”. Microbes and Infection 2.9 (July 2000), pp. 1051–1060. doi: 10.1016/S1286-4579(00)01259-4.

[38] Antonio Mallia, Michał Siedlaczek, and Torsten Suel. “An Experimental Study of Index Compression and DAAT Query Processing Methods”. Advances in Information Retrieval. Ed. by Leif Azzopardi et al. Lecture Notes in Computer Science. Springer International Publishing, 2019, pp. 353–368. isbn: 978-3-030-15712-8.

[39] Marine Le Morvan and Jean-Philippe Vert. “WHInter: A Working Set Algorithm for High-dimensional Sparse Second Order Interaction Models”. Proceedings of the 35th International Conference on Machine Learning. PMLR, July 2018, pp. 3635–3644.

[40] Ankita Nag and Sarika Mehra. “A Major Facilitator Superfamily (MFS) Efflux Pump, SCO4121, from Streptomyces Coelicolor with Roles in Multidrug Resistance and Oxidative Stress Tolerance and Its Regulation by a MarR Regulator”. Applied and Environmental Microbiology 87.7 (), e02238–20. doi: 10.1128/AEM.02238-20.

[41] Ana Ortega-Molina et al. “The Histone Lysine Methyltransferase KMT2D Sustains a Gene Expression Program That Represses B Cell Lymphoma Development”. Nature Medicine 21.10 (Oct. 2015), pp. 1199–1208. doi: 10.1038/nm.3943.

[42] Jakub Otwinowski and Ilya Nemenman. “Genotype to Phenotype Mapping and the Fitness Landscape of the E.Coli Lac Promoter”. PLoS ONE 8.5 (May 2013). Ed. by Andrew J. Yates, e61570. doi: 10.1371/journal.pone.0061570.

[43] Preeti Pachori, Ragini Gothalwal, and Puneet Gandhi. “Emergence of Antibiotic Resistance Pseudomonas Aeruginosa in Intensive Care Unit; a Critical Review”. Genes & Diseases 6.2 (June 2019), pp. 109–119. doi: 10.1016/j.gendis.2019.04.001.

[44] Zheng Pang et al. “Antibiotic Resistance in Pseudomonas Aeruginosa: Mechanisms and Alternative Therapeutic Strategies”. Biotechnology Advances 37.1 (Jan. 2019), pp. 177–192. doi: 10.1016/j.biotechadv.2018.11.013.

[45] powturbo. Powturbo/TurboPFor-Integer-Compression. July 2020.

[46] Pauli Rämö et al. “Simultaneous Analysis of Large-Scale RNAi Screens for Pathogen Entry”. BMC Genomics 15.1 (Dec. 2014), p. 1162. doi: 10.1186/1471-2164-15-1162.

[47] Kay A. Ramsay et al. “Genomic and Phenotypic Comparison of Environmental and Patient-Derived Isolates of Pseudomonas Aeruginosa Suggest That Antimicrobial Resistance Is Rare within the Environment”. Journal of Medical Microbiology 68.11 (Nov. 2019), pp. 1591–1595. doi: 10.1099/jmm.0.001085.

[48] Attika Rehman, Wayne M. Patrick, and Iain L. Lamont. “Mechanisms of Ciprofloxacin Resistance in Pseudomonas Aeruginosa: New Approaches to an Old Problem”. Journal of Medical Microbiology, 68.1 (2019), pp. 1–10. doi: 10.1099/jmm.0.000873.

[49] Attika Rehman et al. “Gene-Gene Interactions Dictate Ciprofloxacin Resistance in Pseudomonas Aeruginosa and Facilitate Prediction of Resistance Phenotype from Genome Sequence Data”. Antimicrobial Agents and Chemotherapy 65.7 (June 2021), e0269620. doi: 10.1128/AAC.02696-20.

[50] Tracey Remmington, Nikki Jahnke, and Christian Harkensee. “Oral Anti-Pseudomonal Antibiotics for Cystic Fibrosis”. Cochrane Database of Systematic Reviews (July 2016). Ed. by Cochrane Cystic Fibrosis and Genetic Disorders Group. doi: 10.1002/14651858.CD005405.pub4.

[51] Ari Robicsek et al. “Fluoroquinolone-Modifying Enzyme: A New Adaptation of a Common Aminoglycoside Acetyltransferase”.Nature Medicine 12.1 (Jan. 2006), pp. 83–88. doi: 10.1038/nm1347.

[52] João V. Rodrigues and Cláudio M. Gomes. “Mechanism of Superoxide and Hydrogen Peroxide Generation by Human Electron-Transfer Flavoprotein and Pathological Variants”. Free Radical Biology and Medicine 53.1 (July 2012), pp. 12–19. doi: 10.1016/j.freeradbiomed.2012.04.016.

[53] Benjamin Schlegel, Rainer Gemulla, and Wolfgang Lehner. “Fast Integer Compression Using SIMD Instructions”. Proceedings of the Sixth International Workshop on Data Management on New Hardware - DaMoN’10. The Sixth International Workshop. Indianapolis, Indiana: ACM Press, 2010, pp. 34–40. isbn: 978-1-4503-0189-3. doi: 10.1145/1869389.1869394.

[54] Fabian Schmich et al. “gespeR: A Statistical Model for Deconvoluting off-Target-Confounded RNA Interference Screens”. Genome Biology 16.1 (Oct. 2015), p. 220. doi: 10.1186/s13059-015-0783-1.

[55] Guomeng Sha et al. “Dynamics and Removal Mechanisms of Antibiotic and Antibiotic Resistance Genes during the Fermentation Process of Spectinomycin Mycelial Dregs: An Integrated Meta-Omics Study”. Journal of Hazardous Materials 421 (Jan. 2022), p. 126822. doi: 10.1016/j.jhazmat.2021.126822.

[56] STRING: Functional Protein Association Networks. https://string-db.org/cgi/about.pl.

[57] Xiaoying Tian, Joshua R Loftus, and Jonathan E Taylor. “Selective Inference with Unknown Variance via the Square-Root Lasso”. Biometrika 105.4 (Dec. 2018), pp. 755–768. doi: 10.1093/biomet/asy045.

[58] Andrew Trotman and Jimmy Lin. “In Vacuo and In Situ Evaluation of SIMD Codecs”. Proceedings of the 21st Australasian Document Computing Symposium. ADCS’16. Caulfield, VIC, Australia: Association for Computing Machinery, Dec. 2016, pp. 1–8. isbn: 978-1-4503-2. doi: 10.1145/3015022.3015023.

[59] UniProt: The Universal Protein Knowledgebase in 2021 — Nucleic Acids Research — Oxford Academic. https://academic.oup.com/nar/article/49/D1/D480/6006196.

[60] Lei Wang et al. “Synergistic Activity of Fosfomycin, Ciprofloxacin, and Gentamicin Against Escherichia Coli and Pseudomonas Aeruginosa Biofilms”. Frontiers in Microbiology 10 (2019), p. 2522. doi: 10.3389/fmicb.2019.02522.

[61] Tong Tong Wu and Kenneth Lange. “Coordinate Descent Algorithms for Lasso Penalized Regression”. The Annals of Applied Statistics 2.1 (Mar. 2008), pp. 224–244. doi: 10.1214/07-AOAS147. arXiv:0803.3876.

[62] Kazuya Yamada et al. “ZHX2 and ZHX3 Repress Cancer Markers in Normal Hepatocytes”. Frontiers in bioscience (Landmark edition) 14 (Jan. 2009), pp. 3724–3732. doi: 10.2741/3483.

[63] Jiyuan Zhang et al. “Disruption of KMT2D Perturbs Germinal Center B Cell Development and Promotes Lymphomagenesis”. Nature Medicine 21.10 (Oct. 2015), pp. 1190–1198. doi: 10.1038/nm.3940.

